# Loss of the ESX-5 secretion locus in *Mycobacterium tuberculosis* reshapes the mycomembrane and enhances ESX-1 substrate secretion

**DOI:** 10.1101/2025.04.11.648440

**Authors:** Benjamin Koleske, Saranathan Rajagopalan, Courtney Schill, Shichun Lun, Catherine Vilchèze, Lahari Das, Manish Gupta, Yazmin B. Martinez-Martinez, William R. Jacobs, William R. Bishai

## Abstract

The ESX-5 secretion system, uniquely found in slow-growing mycobacteria, is predicted to secrete over 150 proteins across the inner membrane of *Mycobacterium tuberculosis* (*M*.*tb*). Although many of these substrates are believed to promote *M*.*tb* virulence, most remain poorly characterized. Here, we use a complete locus deletion strain of ESX-5 in *M*.*tb* to examine the molecular changes caused by a broad loss in ESX-5 secretory substrates. We confirmed the selective loss of PE/PPE proteins secreted by ESX-5 into both the culture filtrate (CF) and outer mycomembrane (OMM) fractions of the *M*.*tb* Δesx5 mutant. In examining other ESX systems, we found that ESX-1 substrate levels were increased in both the CF and OMM fractions of the Δesx5 mutant. Conversely, the ESX-3 locus was transcriptionally repressed upon ESX-5 deletion. We noted that the Δesx5 mutant had altered morphology in the form of wrinkled distortions of the bacterial surface. Likewise, we identified increased susceptibility of the Δesx5 mutant to a variety of large (molecular weight >550 g/mol) antimicrobial compounds, suggesting that an intact ESX-5 system is required for *M*.*tb* to exclude such molecules. Our findings suggest that removing the ESX-5 system from *M*.*tb* fundamentally alters the properties of the mycobacterial OMM and impacts the expression and secretion activity of other ESX systems.

**Significance Statement:** *Mycobacterium tuberculosis* (*M*.*tb*) uses the ESX-5 secretion system to export numerous proteins that shape host-pathogen interactions. Here, we found that deleting ESX-5 from *M*.*tb* not only prevented the secretion of many ESX-5 substrates but also impacted other ESX systems. The *M*.*tb* Δesx5 mutant had increased ESX-1 substrate secretion but reduced ESX-3 expression. In addition, the *M*.*tb* Δesx5 mutant displayed altered cell surface morphology and increased vulnerability to large antibiotic drugs, suggesting a critical role for ESX-5 for maintaining outer membrane integrity. These findings highlight ESX-5 as a central modulator of secretion and cell envelope composition with implications for drug targeting and vaccine development.

## Introduction

The staggering global burden of *Mycobacterium tuberculosis* (*M*.*tb*) reflects the challenges of eradicating an organism naturally resistant to many antimicrobials and host immune defenses (1, 2). A key mediator of this intrinsic resistance is the *M*.*tb* cell wall, which features a distinctive outer mycomembrane (OMM) composed of large waxy mycolic acid lipids (3, 4). However, the adaptive advantage conferred by this robust barrier is met with a corresponding disadvantage, as *M*.*tb* must also import nutrients and export its secretory effectors across this poorly permeable layer. Mycobacteria use their type VII secretion systems (T7SSs), the ESX systems, to transport proteins into and across the OMM via a bipartite secretion motif (5, 6). Among the five T7SSs in *M*.*tb*, ESX-1 is critical for phagosomal rupture, and its loss from the *M. bovis* BCG vaccine strain is a major driver of attenuation (7, 8). The ESX-3 system facilitates the acquisition of metal ions, principally iron and zinc, although it has also been recently implicated in calcium uptake (9–11). Roles for ESX-2 and ESX-4 remain unclear, although recent work has suggested that they may support the functions of ESX-1 or ESX-3 (12–14).

ESX-5 is the youngest of the *M*.*tb* T7SSs evolutionarily, yet it has by far the largest diversity of putative secretory substrates (15). The duplication of ESX-5 from ESX-2 corresponds with both the transition to the slow-growing lifestyle employed by many pathogenic mycobacteria and an extensive expansion of T7SS substrates, chiefly the PE (Pro-Glu) and PPE (Pro-Pro-Glu) families (15). These families, named for motifs in their conserved N-terminal secretion domains, comprise 168 members in *M*.*tb* compared to 11 members in the non-pathogenic *Mycobacterium smegmatis* (16–18). Most PE/PPE proteins are predicted to be secreted across the cytoplasmic membrane by ESX-5, and indeed many localize to the OMM (19–21). Despite their abundance in the genome, most PE/PPE proteins lack a known function and can be individually knocked out by transposon mutagenesis with minimal apparent *in vitro* fitness cost to the bacterium (22, 23). Notably, a number of PE/PPE proteins are highly induced during animal infection, suggesting that they may contribute to pathogenesis (24). As secretory system substrates, the numerous PE/PPE proteins also represent a large repertoire of possible epitopes exposed to the host immune system (20, 25, 26). The continued maintenance of these large substrate families in an otherwise reduced pathogen genome highlights the importance of these proteins on the *M*.*tb* life cycle (15, 16, 27).

Mutation of the ESX-5 machinery may therefore alter the localization of upwards of 150 *M*.*tb* proteins, many of which have been proposed to function in host-pathogen interactions and immune evasion (25, 28). Such a substantial shift in the culture filtrate (CF) or OMM compartments may also create vulnerabilities in *M*.*tb*, as several PE/PPE substrates of ESX-5 are known to facilitate small molecule uptake (29). Hence, it is possible that a complete ESX-5 deletion strain would possess altered properties beyond the mere loss of ESX-5 substrate secretion. Prior work on disrupting the ESX-5 locus in different *M*.*tb* strains, as well as *M. marinum*, has shown a range of effects, from non-viability to attenuation and even hypervirulence (20, 30–34). A complete deletion of the ESX-5 locus has been achieved in *M*.*tb* CDC1551; while this deletion causes attenuation in immunocompromised mice, there has not yet been a protein-level investigation of the fate of ESX-5 substrates in such a strain (34). This may be due in part to the high similarity between PE/PPE proteins, which makes them challenging to disambiguate by mass spectrometry. Of particular difficulty are the PE_PGRS proteins, a modern subfamily of PE proteins with a polymorphic GC-rich repeat sequence (PGRS) domain composed of roughly two-thirds glycine residues. We were encouraged by the extensive work to characterize *PPE38* and *PPE71* as necessary for PE_PGRS export, as this work represented solid proof that large shifts in ESX-5 substrate compartmentalization can be assessed experimentally (26, 35, 36). Notably, disrupting secretion of the PE_PGRS proteins en bloc helped overcome the risk of possible cross-complementation between the highly similar PE_PGRS family members. We believe that directly ablating the ESX-5 system may similarly help overcome potential functional redundancy that may conceal the functions of single *PE/PPE* gene mutants.

In this study, we used untargeted mass spectrometry to determine differences in the proteomes of the CF and OMM fractions of an *M*.*tb* Δesx5 mutant compared to the parental CDC1551 WT strain. We observed widespread loss in diverse PE/PPE proteins, including several with no prior experimental evidence of ESX-5 secretion. Despite this loss, *PE_PGRS* gene transcription was upregulated despite an absence of secretion, possibly as a compensatory mechanism. Intriguingly, multiple ESX-1 substrates were increased in both the CF and OMM fractions, while expression of ESX-3 was decreased at the transcript level. In addition, the Δesx5 mutant exhibited altered cell surface morphology by scanning electron microscopy and was more susceptible to a wide range of antibiotic drugs, particularly molecules with higher molecular weights. In summary, using a novel full-locus deletion of ESX-5, we identify new members of the ESX-5 ‘secretome’ and demonstrate changes in the ESX-1 and ESX-3 systems as well as cell wall morphology and drug susceptibility.

## Results

### Proteomic profiling of *M*.*tb* Δesx5 secreted fractions

The ESX-5 locus of *M*.*tb* encompasses 17 genes from *eccB5* (Rv1782) to *eccA5* (Rv1798), which collectively span just under 21 kilobases (16). We used specialized transduction to replace the ESX-5 locus of *M*.*tb* CDC1551 with a hygromycin resistance cassette and designated this strain “Δesx5” (37). Specifically, we constructed an allelic exchange substrate mycobacteriophage with a central hygromycin resistance cassette and a *sacB* counter-selection marker flanked by ~1 kilobase of *M*.*tb* genomic DNA from immediately upstream and downstream of the ESX-5 operon. Upon transduction, we selected colonies on hygromycin and further screened by PCR to identify deletion mutants (**Fig. S1**).

Because the ESX-5 system secretes proteins into the OMM as well as the CF, we sought a workflow to isolate the OMM fraction while minimizing contamination from bacterial cytoplasmic proteins. To this end, we used n-octyl β-D-glucoside (OBG), a mild non-ionic detergent that has been shown to solubilize the OMM without disrupting mycobacterial cell integrity (38, 39). We confirmed that incubation with OBG did not cause appreciable cell lysis, as there was no significant difference in viability between *M*.*tb* WT or Δesx5 cultures incubated with or without OBG as assessed by CFU counts (**Fig. S2**). Additionally, we did not detect any bacterial chromosomal DNA in the extracted fraction following OBG treatment, suggesting again that the bacteria did not lyse under this treatment. To promote the formation of a robust OMM, we omitted the detergent from a previously described recipe for a mildly acidic (pH 6.5) 7H9-based induction medium (40, 41). This medium also contains propionate, which promotes the production of branched lipids, a key component of the OMM (40, 42).

Following induction, we isolated and filtered the CF before directly incubating intact mycobacteria with OBG to extract the OMM fraction (**Fig. 1A**). We subjected CF and OMM fractions to untargeted mass spectrometry and observed clear separation between WT and Δesx5 strains in both CF and OMM fractions by principal component analysis (**Fig. 1B–C**). This confirmed that the Δesx5 strain showed consistent, detectable differences in both CF and OMM proteins compared to WT.

**Figure 1:**
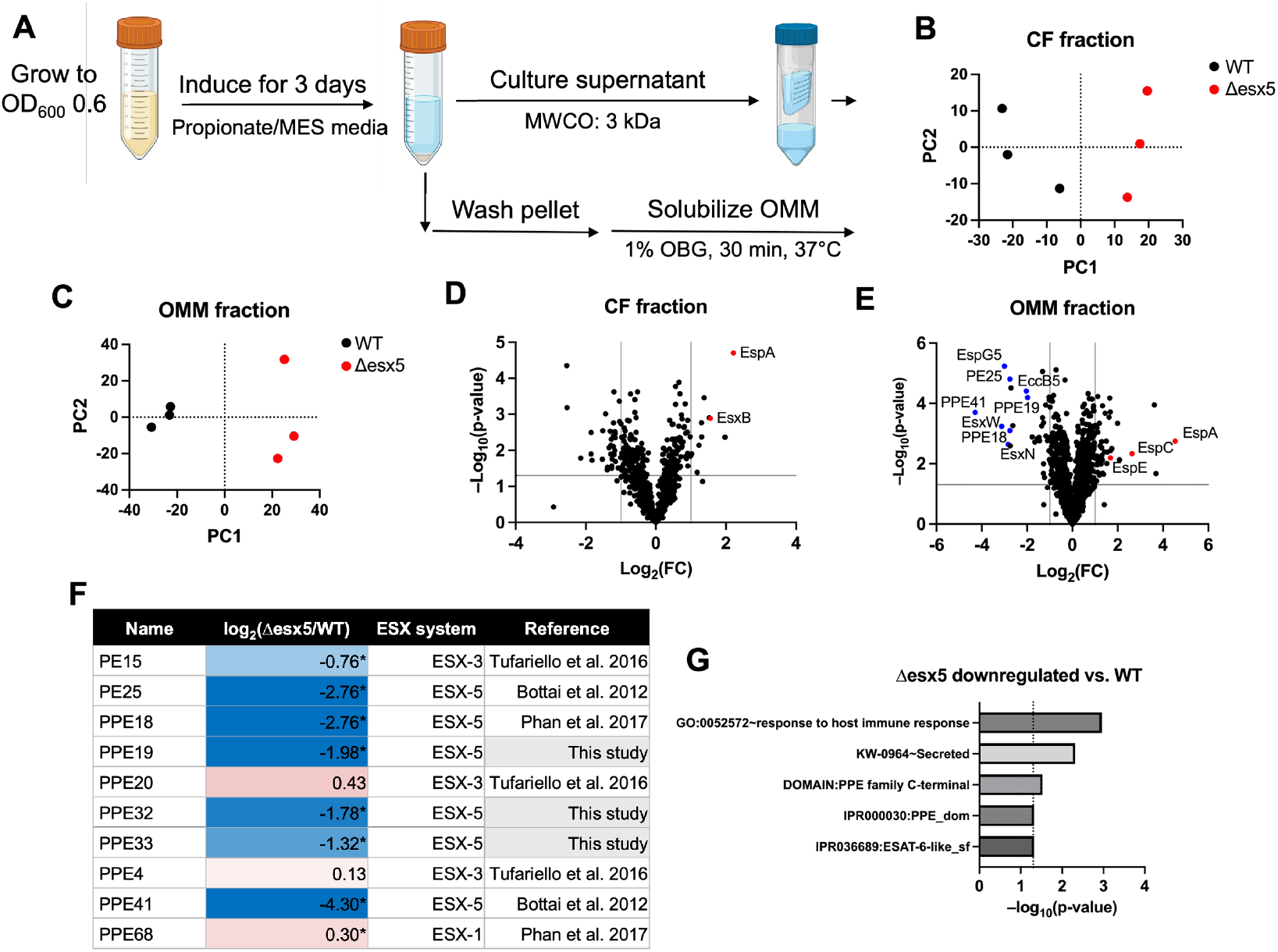
*M*.*tb* ESX-5 secretes a plethora of PE/PPE proteins predominantly into the outer cell wall. (A) Workflow for extraction of soluble secreted and outer mycomembrane (OMM) proteins from *M*.*tb* for mass spectrometry. Cultures were grown to a mid-log OD_600_ of 0.6 and induced in propionate/MES induction media for 3 days. Culture supernatant was harvested by centrifugal concentration through a 3 kilodalton (kDa) molecular-weight cutoff (MWCO) filter to generate culture filtrate (CF). The cell pellet was washed and incubated in 1% octyl glucoside (OBG) in TBS for 30 min at 37°C, and the soluble OMM fraction was harvested for analysis. (B–C) Principal component (PC) plots derived from proteins detected by mass spectrometry in the CF (B) and OMM (C) fractions of WT and Δesx5 *M*.*tb* strains (n=3). (D–E) Volcano plots of proteins identified by mass spectrometry in the CF (D) and OMM (E) fractions of the Δesx5 strain normalized to the WT strain. Grey lines indicate thresholds of ±1 log_2_(fold-change) (vertical) and p < 0.05 (horizontal). Select downregulated PE/PPE proteins and ESX-5-associated proteins are colored in blue, while upregulated ESX-1-associated proteins are colored in red. (F) List of PE/PPE family proteins detected in the OMM fraction, alongside their abundance in the Δesx5 strain compared to WT (log_2_(fold-change)) and the ESX system responsible for secreting each protein. References are provided for experimentally confirmed secretion by the respective ESX system. If a protein had only evolutionary or computational prediction of secretion by ESX-5, it is attributed to ‘this study’. (*: p < 0.05.) (G) Significant functional annotation terms for proteins downregulated by at least 2-fold (p < 0.05) in the OMM fraction of the Δesx5 strain compared to WT.

### *M*.*tb* Δesx5 fails to secrete many PE/PPE proteins into the OMM

Next, we examined the individual proteins detected in each secretory fraction, with a focus on putative T7SS substrates. Most PE/PPE proteins are known or predicted to be ESX-5 substrates, though a handful are secreted by other T7SSs, mainly ESX-3 (15, 43). We did not detect any PE/PPE proteins in the CF fractions of either WT or the Δesx5 mutant (**Fig. 1D, Dataset S1**). Conversely, PPE proteins were abundant in the OMM fraction, as were several PE proteins and small Esx-family proteins (**Fig. 1E, Dataset S2**). We found that over half of the PE/PPE proteins we detected had significantly decreased abundance in the Δesx5 mutant compared to WT (**Fig. 1F**). As would be predicted, the PE/PPE proteins that did not decrease in the Δesx5 mutant are all known to be secreted by other ESX secretion systems, confirming that the Δesx5 mutant has a selective secretion defect for ESX-5 substrates (20, 43, 44). Functional annotation clustering via the Database for Annotation, Visualization, and Integrated Discovery (DAVID) uncovered a cluster of terms corresponding to host immune responsiveness and PPE proteins that were significantly downregulated in the OMM fraction of the Δesx5 mutant compared to WT (**Fig. 1G**). Thus, the Δesx5 mutant has impaired secretion of a plethora of PPE proteins into the mycobacterial OMM yet continues to secrete ESX-1 and ESX-3 substrates.

### The *M*.*tb* Δesx5 mutant has increased secretion of ESX-1 substrates

Initially, we hypothesized that the greatest differences between the WT and Δesx5 mutant secretory fractions would be the loss of putative ESX-5 substrates. Surprisingly, we also found that a number of ESX-1 system substrates were upregulated in the CF and OMM fractions of the Δesx5 mutant (**Fig. 1D–E**). A more comprehensive examination of ESX-1 system substrates demonstrated significant upregulation of several more such proteins in both secretory fractions of the Δesx5 mutant (**Fig. 2A–B**). Indeed, EspA had the single greatest increase in either fraction, as it was nearly 5-fold more abundant in the CF fraction and 23-fold more abundant in the OMM fraction of the Δesx5 mutant compared to WT.

**Figure 2:**
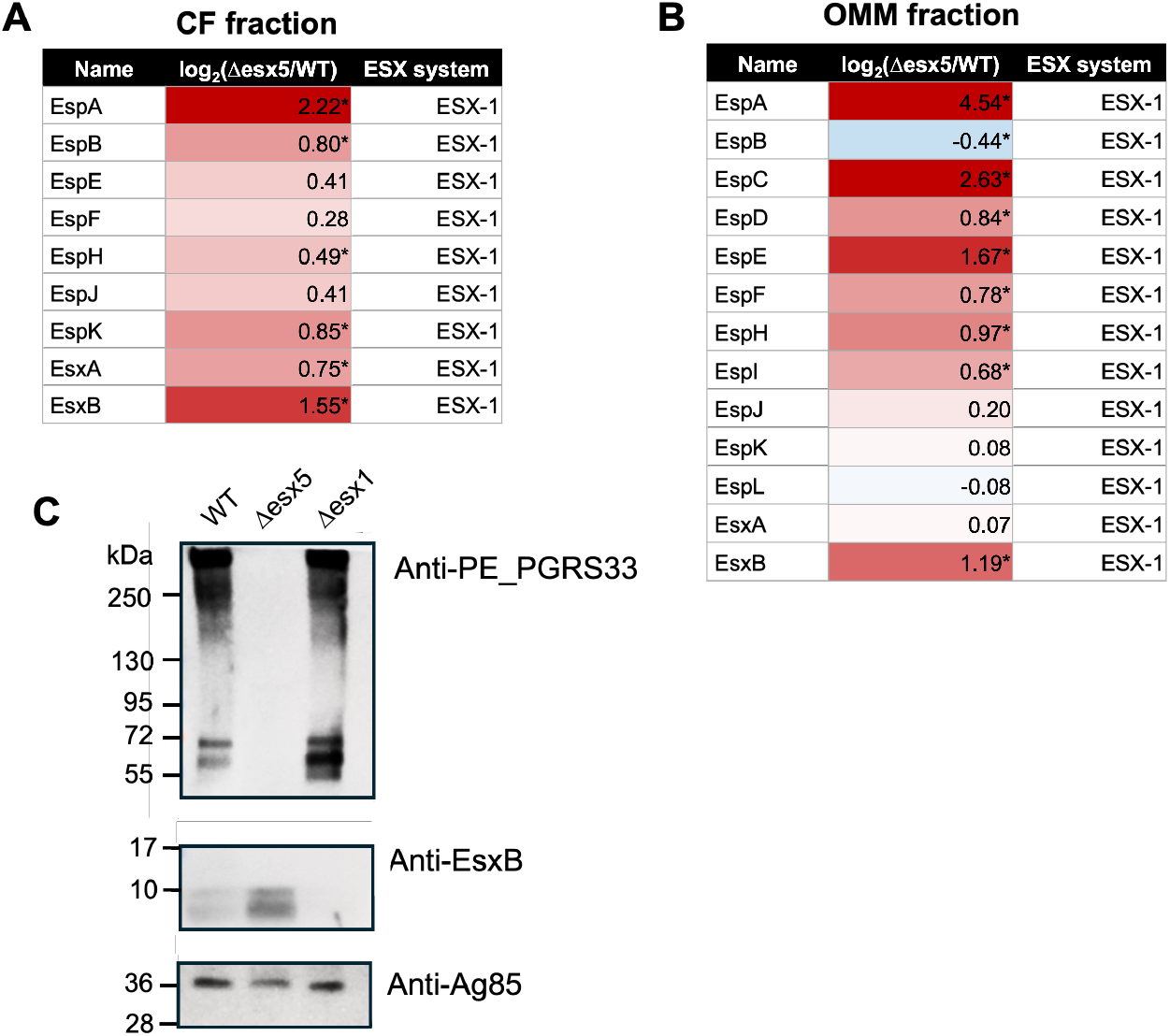
The *M*.*tb* Δesx5 mutant upregulates the secretion of several ESX-1 substrates. (A–B) List of ESX-1 substrates found in the culture filtrate (CF) (A) and outer mycomembrane (OMM) (B) fractions and their enrichment (log_2_(fold-change) in the Δesx5 strain. (*: p < 0.05.) (C) Western blots of OMM fractions from WT, Δesx5, and Δesx1 *M*.*tb* strains for PE_PGRS33 (top), EsxB (middle), and Ag85 (bottom). (kDa: kilodaltons.)

To validate this finding of increased ESX-1 substrate secretion in the Δesx5 mutant, we subjected OMM fractions of the *M*.*tb* WT and Δesx5 strains to Western blotting. As a control, we used the OMM fraction from a Δesx1 mutant strain, which would be expected to lose ESX-1 secretion but preserve ESX-5 secretion. The Δesx5 strain failed to secrete PE_PGRS family proteins into the OMM, consistent with the known role of ESX-5 in PE_PGRS secretion (**Fig. 2C**, top) (19, 35). Conversely, the Δesx5 mutant secreted considerably more EsxB, a stable and highly immunogenic ESX-1 substrate (45), than WT (**Fig. 2C**, middle). Neither the Δesx5 nor the Δesx1 mutant showed a difference from WT in the secretion of Antigen 85 (Ag85), a twin-arginine translocation system substrate used a control (**Fig. 2C**, bottom) (46). Hence, the increase in ESX-1 substrate secretion seen in the Δesx5 strain does not reflect a nonspecific upregulation of all remaining *M*.*tb* secretion systems upon loss of ESX-5.

### The *M*.*tb* Δesx5 mutant transcriptionally upregulates *PE_PGRS* genes and downregulates the ESX-3 locus

To establish that the defect in ESX-5 substrate secretion seen in the Δesx5 mutant was due to impaired protein localization rather than transcriptional repression, we performed RNA-seq on transcripts from WT and Δesx5 strains. The *PE_PGRS* family of ESX-5 substrates stood out as being strongly upregulated in the Δesx5 mutant compared to WT (**Fig. 3A**). Among the 61 *PE_PGRS* transcripts we detected, 32 (52%) showed a significant increase of at least 2-fold in the Δesx5 mutant. No *PE_PGRS* transcripts were decreased by 2-fold or greater. As a result, functional analysis via DAVID identified a highly enriched *PE_PGRS* cluster among genes significantly upregulated in the Δesx5 mutant (**Fig. 3B**). The inability of the Δesx5 strain to export PE_PGRS proteins is therefore not due to poor transcription of these genes. Instead, the broad upregulation of *PE_PGRS* transcripts may reflect an attempt by the Δesx5 mutant to compensate for a deficiency in properly localized PE_PGRS proteins.

**Figure 3:**
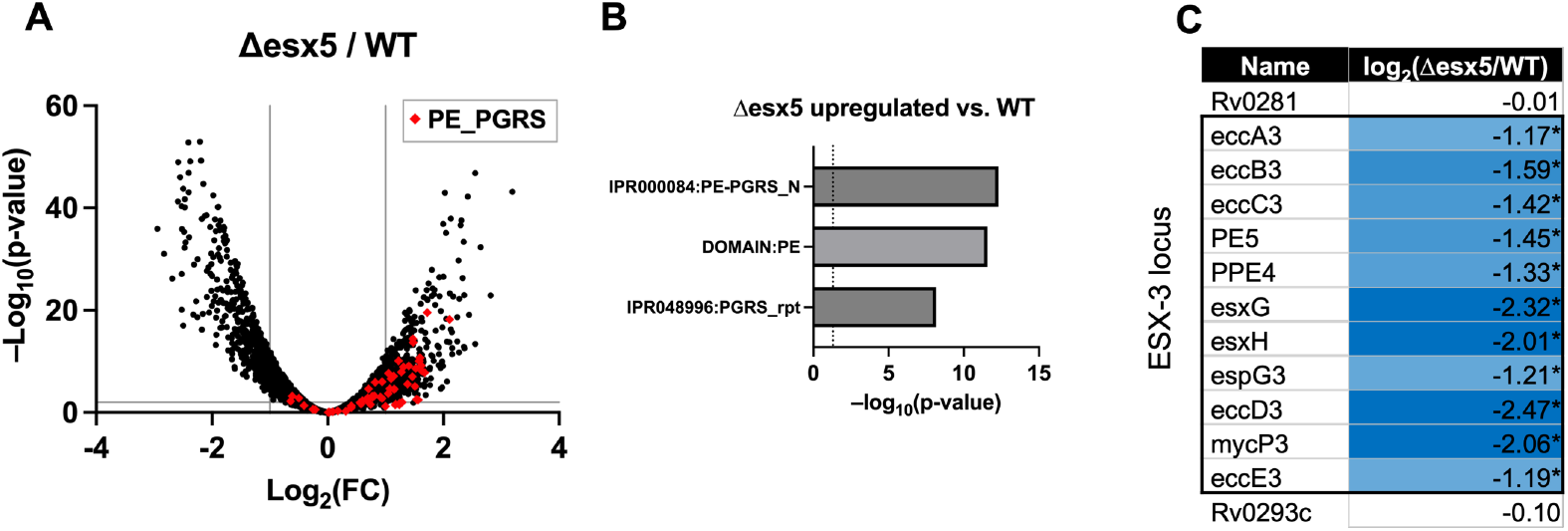
The *M*.*tb* Δesx5 mutant transcriptionally upregulates PE_PGRS genes and downregulates the ESX-3 locus. (A) Volcano plot of transcripts quantified in the Δesx5 strain normalized to WT. Grey lines indicate thresholds of ±1 log_2_(fold-change) (vertical) and p < 0.01 (horizontal). PE_PGRS family genes are indicated with red diamonds. (B) Top significant functional annotation terms for transcripts upregulated by at least 2-fold (p < 0.01) in the Δesx5 strain compared to WT. (C) Transcriptional expression of ESX-3 locus genes in the Δesx5 strain compared to WT (log_2_(fold-change)). Flanking genes on either side of the ESX-3 locus are provided for contrast. (*: p < 0.05.)

In light of the apparent increase in ESX-1 substrate secretion in the Δesx5 strain, we were interested in whether transcription of the ESX-1 core machinery or ESX-1 substrate loci had increased. Neither the ESX-1 system nor its *esp*-family substrates were transcriptionally upregulated in the Δesx5 mutant; in fact, the transcripts for certain ESX-1 substrates (e.g., the *espACD* locus) were downregulated (**Dataset S3**). We noticed, however, that the ESX-3 core machinery locus was significantly repressed at the transcriptional level in the Δesx5 mutant (**Fig. 3C**). Several known ESX-3 substrates, such as the PE15/PPE20 pair, were also downregulated in the Δesx5 mutant, although this was not consistent across all known and predicted ESX-3 substrates (**Dataset S3**) (43).

### The *M*.*tb* Δesx5 mutant has altered bacterial surface morphology

During our proteomic experiments, we observed that the Δesx5 mutant displayed different properties than WT and Δesx1 mutant bacteria. The WT and Δesx1 strains formed clumps and waxy adhesions at the air-liquid interface of the culture tube when grown in induction medium, which is the expected behavior for *M*.*tb* in a medium without detergent and containing propionate to promote branched lipid production (41, 42).

Unlike these strains, the Δesx5 mutant formed minimal clumps and instead grew largely planktonically under the same conditions. Hence, we wondered whether the surface properties of Δesx5 mutant bacteria differed from the WT and Δesx1 mutant strains. We adhered WT, Δesx5, and Δesx1 strains to coverslips and subjected them to scanning electron microscopy (SEM) to image cell morphology. In rich media, WT and Δesx1 mutant bacteria had a smooth, rounded surface consistent with normal mycobacterial morphology, but the Δesx5 mutant had pronounced wrinkles spanning up to the full length of the cells (**Fig. 4A–C**). Incubating the bacteria in the induction medium further exacerbated the wrinkled surface texture of the Δesx5 mutant but did not affect the WT strain (**Fig. 4D–E**). We subjected WT and Δesx5 mutant cultures to ^14^C-propionate labeling and identified diminished levels of apolar OMM lipid species in the Δesx5 mutant (**Fig. S4**). We also measured the lengths of WT and Δesx5 mutant bacteria from the SEM images and found that Δesx5 mutant cells (mean 1.92 µm) were significantly shorter than WT cells (mean 2.31 µm) (**Fig. 4F**).

**Figure 4:**
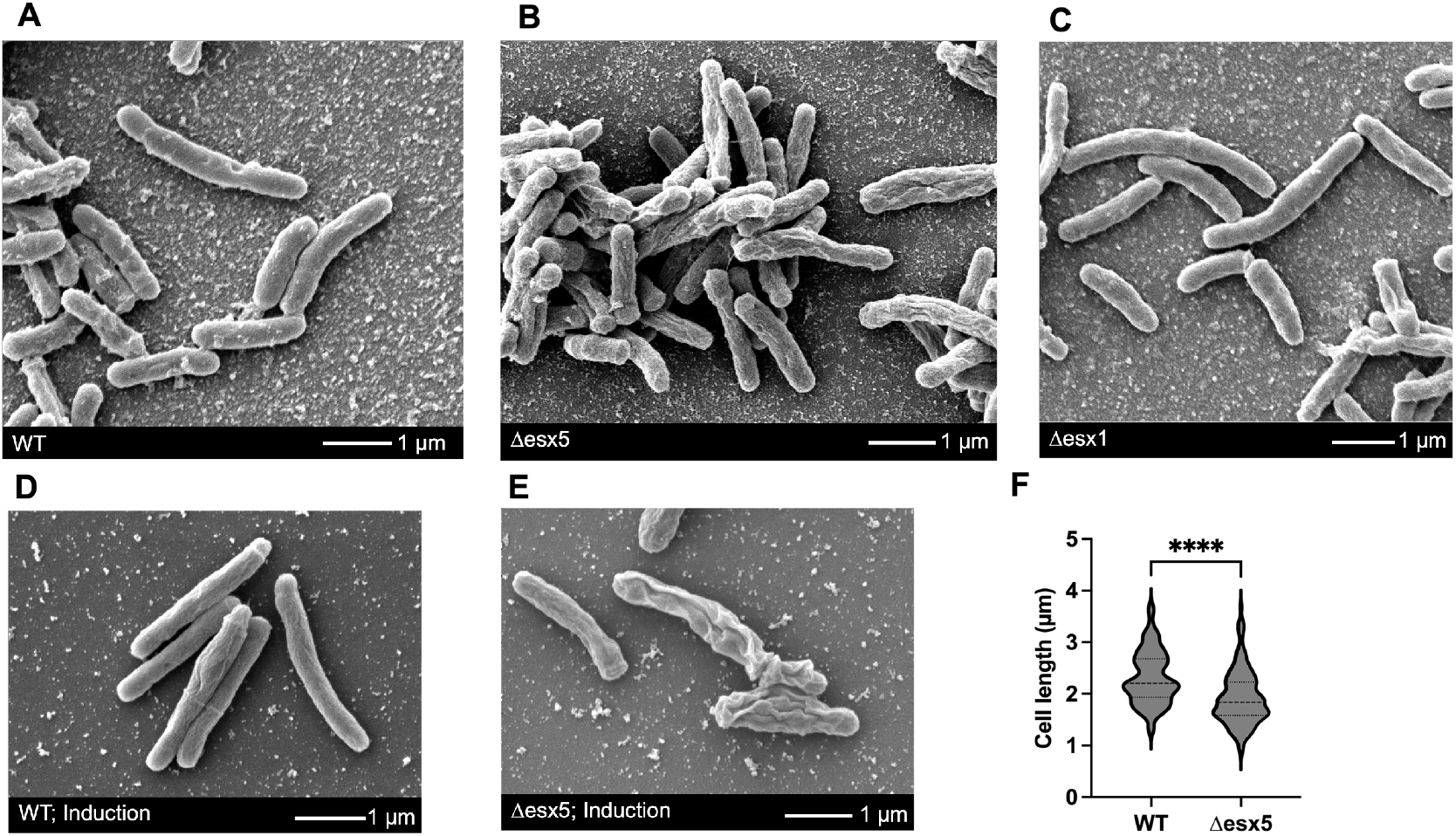
The *M*.*tb* Δesx5 mutant displays an altered cell morphology. (A–C) Scanning electron microscopy images of WT (A), Δesx5 (B), and Δesx1 (C) *M*.*tb* strains. Bacteria were grown on coverslips for 48 hrs in 50:50 complete 7H9 : rich YM media with 6 mM DTT to promote adhesion. (D–E) Scanning electron microscopy images of WT (D) and Δesx5 (E) *M*.*tb* strains. Bacteria were grown on coverslips for 48 hrs in propionate/MES induction media with 6 mM DTT to promote adhesion. (F) Cell length distributions for WT and Δesx5 *M*.*tb* strains grow in 50:50 7H9:YM media, as quantified from SEM images. (n>100 each, ****: p<0.0001 by unpaired two-tailed t test.)

In addition to SEM, we plated the WT, Δesx1, and Δesx5 strains on 7H11 agar to examine colony morphology. At six weeks after seeding, WT and Δesx1 mutant strains displayed rough, irregularly shaped waxy colonies typical of *M*.*tb* (**Fig. S3A–B**). At the same time point, the Δesx5 mutant colonies were smooth, round, and smaller overall (**Fig. S3C**). Thus, the Δesx5 mutant displayed an altered growth morphology across multiple culture modalities.

### The *M*.*tb* Δesx5 mutant is more susceptible to many antimicrobials, particularly those with higher molecular weights

Due to the differences in surface morphology and mycomembrane composition, we wondered whether the Δesx5 mutant would show differential antibiotic susceptibility. In particular, the mycobacterial OMM is believed to impair the uptake of large or polar antimicrobial compounds (1). Using a microplate-based alamarBlue assay (47, 48), we determined the minimum inhibitory concentration (MIC) values for a diverse range of antimicrobial drugs against *M*.*tb* WT and Δesx5 mutant strains. Of the 20 drugs tested, we observed that 14 showed an MIC at least 2-fold lower for Δesx5 compared to WT, including 8 drugs that showed an MIC at least 8-fold lower (**Table 1**). No drugs had a higher MIC for the Δesx5 mutant. Upon examining the properties of these molecules, we found that the drugs with the greatest change in MIC were among those with the highest molecular weights. Indeed, among the antimicrobials with molecular weights of at least 550 g/mol, all drugs except the three aminoglycosides (streptomycin, kanamycin, and amikacin) had a ≥8-fold decrease in the MIC for the Δesx5 mutant compared to WT. The greatest decreases in MIC were seen with vancomycin (512-fold lower in the Δesx5 mutant), azithromycin (128-fold lower), and the two tested rifamycins, rifampin and rifapentine (each 32-fold lower). Hence, Δesx5 has broadly increased susceptibility to a range of antibiotics and appears particularly sensitive to larger compounds. This may be a consequence of altered cell wall or cell surface properties.

**Table 1:**
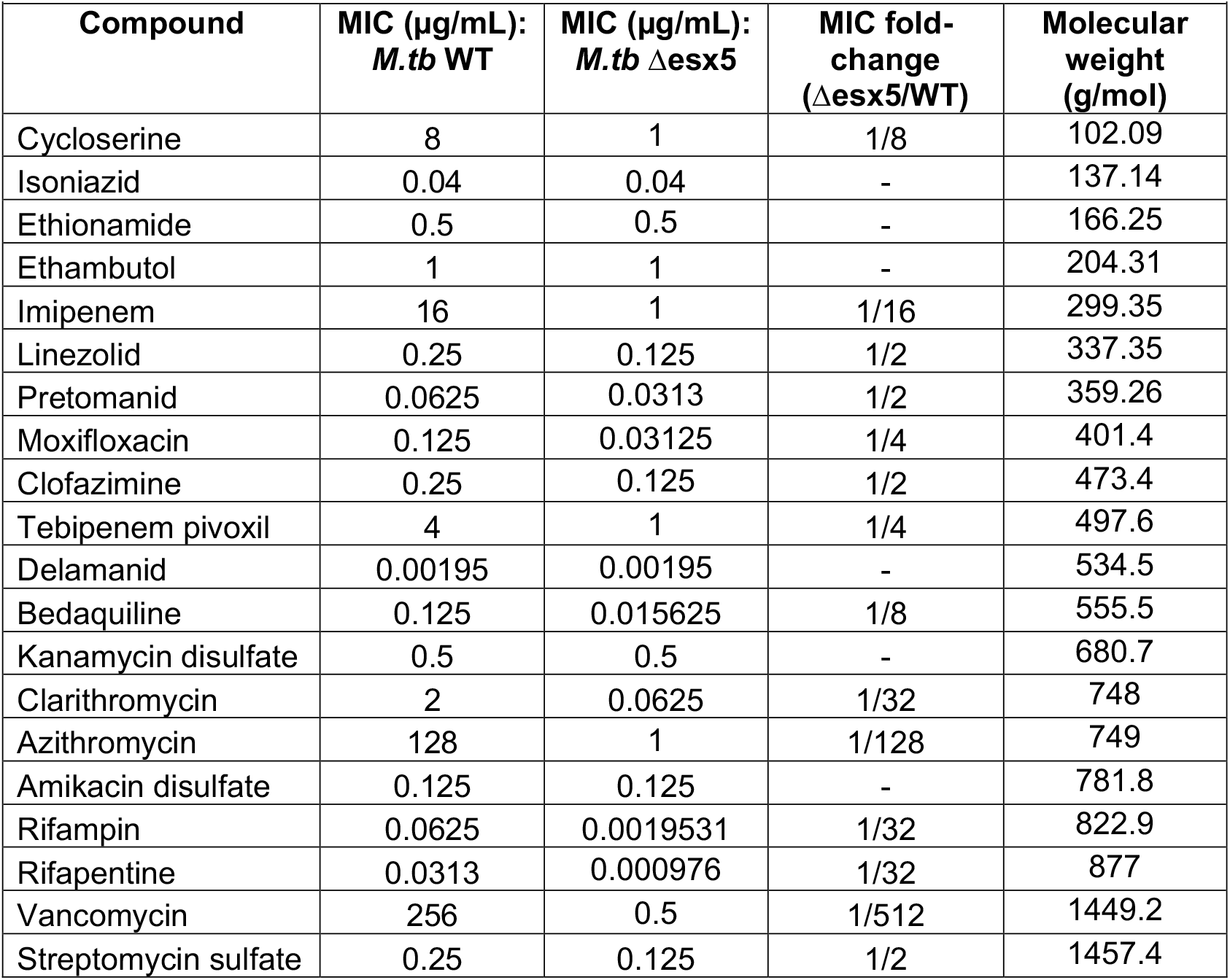
MIC values of antimicrobial drugs against *M*.*tb* WT and Δesx5 strains.

| Table 1: MIC values of antimicrobial drugs against *M.tb* WT and Δesx5 strains. | | | | |
| --- | --- | --- | --- | --- |
| Compound | MIC (μg/mL):<br>*M.tb* WT | MIC (μg/mL):<br>*M.tb* Δesx5 | MIC fold-change<br>(Δesx5/WT) | Molecular<br>weight<br>(g/mol) |
| Cycloserine | 8 | 1 | 1/8 | 102.09 |
| Isoniazid | 0.04 | 0.04 | - | 137.14 |
| Ethionamide | 0.5 | 0.5 | - | 166.25 |
| Ethambutol | 1 | 1 | - | 204.31 |
| Imipenem | 16 | 1 | 1/16 | 299.35 |
| Linezolid | 0.25 | 0.125 | 1/2 | 337.35 |
| Pretomanid | 0.0625 | 0.0313 | 1/2 | 359.26 |
| Moxifloxacin | 0.125 | 0.03125 | 1/4 | 401.4 |
| Clofazimine | 0.25 | 0.125 | 1/2 | 473.4 |
| Tebipenem pivoxil | 4 | 1 | 1/4 | 497.6 |
| Delamanid | 0.00195 | 0.00195 | - | 534.5 |
| Bedaquiline | 0.125 | 0.015625 | 1/8 | 555.5 |
| Kanamycin disulfate | 0.5 | 0.5 | - | 680.7 |
| Clarithromycin | 2 | 0.0625 | 1/32 | 748 |
| Azithromycin | 128 | 1 | 1/128 | 749 |
| Amikacin disulfate | 0.125 | 0.125 | - | 781.8 |
| Rifampin | 0.0625 | 0.0019531 | 1/32 | 822.9 |
| Rifapentine | 0.0313 | 0.000976 | 1/32 | 877 |
| Vancomycin | 256 | 0.5 | 1/512 | 1449.2 |
| Streptomycin sulfate | 0.25 | 0.125 | 1/2 | 1457.4 |

## Discussion

The ESX-5 secretion system and its greatly expanded PE/PPE substrate repertoire are hallmarks of slow-growing pathogenic mycobacteria and likely play important roles during host infection. Here, we present a molecular characterization of a novel, full-locus deletion of the ESX-5 system in *M*.*tb*. Using this *M*.*tb* Δesx5 mutant, we have demonstrated the ESX-5 dependence of several OMM-associated PPE proteins.

Notably, we detected increased secretion of ESX-1 substrates, including Esp-family proteins and EsxB, into the CF and OMM fractions of the Δesx5 mutant. Corresponding to the lack of ESX-5 substrate secretion, we noted a loss of PE_PGRS proteins in the OMM fraction, which was accompanied by a compensatory increase in *PE_PGRS* family transcripts. Unexpectedly, we found transcriptional downregulation of the ESX-3 locus in the Δesx5 mutant. Using SEM, we observed wrinkled cell surface distortions in the Δesx5 mutant strain consistent with altered OMM behavior. Lastly, we detected greater susceptibility of the Δesx5 mutant to a range of antimicrobial compounds, particularly those with a high molecular weight.

In this study, we adapted a previously described culture medium that incorporates propionate into a 7H9 base alongside mild acid stress, which partly mimics the host phagosome (40). Using these induction conditions, we definitively identified 27 ESX system substrates in our secretory fractions by mass spectrometry, including 2 PE, 8 PPE, 6 Esx, and 11 Esp family proteins. It is not surprising that we achieved a partial list of the complete *M*.*tb* T7SS substrate repertoire, as these protein families have long been considered challenging to study. PE/PPE proteins and small Esx proteins have high similarity between family members and few basic amino acids for trypsin cleavage (49, 50). Due to these properties, these proteins may not generate sufficient unique peptides for detection by mass spectrometry. Nonetheless, we identified three PPE proteins, PPE19, PPE32, and PPE33, that had previously lacked experimental confirmation as ESX-5 substrates. In this study, we focused on a single culture condition that used acid stress as an analog of the host phagosome. However, a single stress condition may not elicit high-level expression of all ESX-5 substrates. Given that several PE/PPE proteins appear to serve as small molecule transporters or host-interacting proteins, we suspect that different family members are inducible under different conditions (29, 50, 51). For example, phosphate starvation has been found to induce expression of the ESX-5 locus and several of its putative substrates (52). It may also be informative to generate a similar Δesx5 deletion in different *M*.*tb* strain backgrounds. In certain backgrounds, including H37Rv, ESX-5 appears to be strictly essential (30), although Δesx5 strains can be generated in CDC1551 as reported here and previously (34). Future work using a more comprehensive range of conditions and strains may allow for the validation of additional ESX-5 substrates at the protein level.

As seen from our proteomic data, the Δesx5 mutant secretes a substantially different set of proteins into both the mycobacterial OMM and the soluble CF compartment. The PE_PGRS proteins are entirely absent from the OMM fraction of the Δesx5 mutant, consistent with their known dependence on ESX-5 for localization (19, 35). This protein family is highly secreted in animal infection and has been proposed to subvert host immunity via the highly repetitive, poorly immunogenic PGRS domain (24, 53, 54).

Interestingly, in lieu of PE_PGRS proteins, the Δesx5 strain demonstrated increased secretion of ESX-1 substrates from two distinct genetic loci. The mechanism for this increase remains unclear, although it is interesting to consider that a partially attenuated pathogen like the Δesx5 mutant may respond to a state of relative stress by upregulating its remaining virulence factors. Several transcriptional regulators of *M*.*tb* virulence genes, including Lsr2 and EspR, regulate both the ESX-1 and ESX-5 loci, allowing the possibility of feedback between these systems (55–57). Additionally, EspA in particular is known to promote *M*.*tb* cell wall integrity, a process with which the Δesx5 mutant likely struggles due to its morphology defects (58, 59). Hence, it remains unclear whether the apparent crosstalk between different ESX systems may be due to direct transcriptional co-regulation or mediated by a central process, such as sensing alterations in OMM composition. Either way, the loss of over 150 cell wall and secretory proteins all but ensures that the Δesx5 mutant will present a highly different set of peptides to the host environment. Hence, we anticipate that a rigorous comparison of the Δesx5 mutant and WT strains in cellular and animal assays may demonstrate a role for ESX-5 system substrates in host immune interactions.

The dramatic wrinkled surface of the Δesx5 strain, as seen by SEM, suggests a considerable change in the structure or composition of the OMM of this strain. This phenotype can be explained by the link between the ESX-5 system and the permeability of the mycomembrane to small molecules (29, 31). Indeed, mutations in several PE/PPE nutrient transporters of ESX-5 or the ESX-5 core machinery have been shown to cause non-viability unless complemented by either detergent treatment or expression of porin transgenes to permeabilize the OMM (29, 31). With a diminished set of such proteins, the OMM of the Δesx5 mutant is likely to be dramatically altered both structurally and functionally from that of WT *M*.*tb*. We hypothesize that the Δesx5 mutant lacks important OMM components and that this impaired architectural integrity leads to greater permeability of the OMM. This interpretation is supported by the reduced MICs for the Δesx5 strain across a broad range of high molecular weight antibiotics, seemingly irrespective of the drugs’ targets. Yet, the Δesx5 mutant appears to retain some ability to exclude the most highly polar molecules, as it is not more susceptible than WT to the positively charged aminoglycosides. Prior work has found that recombinant expression of the *M. smegmatis* major porin *mspA* gene into *M*.*tb* increased both nutrient uptake and susceptibility to numerous antibiotics, establishing a link between these properties and OMM permeability (60). The perturbations in the OMM may also explain the unintuitive repression of the ESX-3 system—which mediates iron, zinc, and calcium ion uptake (9–11)—in the Δesx5 mutant. While the ESX-3 system is essential under ordinary conditions, an increase in OMM permeability may allow siderophores to traverse the cell wall more readily without requiring ESX-3 activity.

Further detailed metabolic profiling of the Δesx5 mutant may uncover other changes needed to sustain this drastic mutation.

Taken together, we have established that deletion of the ESX-5 T7SS in *M*.*tb* causes loss of numerous PE/PPE and Esx-family substrates from the mycobacterial OMM. Beyond ESX-5, we observed changes in the two other major *M*.*tb* T7SSs, with ESX-1 hypersecretion and ESX-3 repression in the Δesx5 mutant. The Δesx5 mutant further showed altered cell surface and colony morphology. Furthermore, the Δesx5 mutant appeared more susceptible to large antimicrobial compounds of moderate or low polarity. Given the vital, pathogen-defining role of ESX-5, we believe that further exploration using our *M*.*tb* Δesx5 strain may demonstrate how this system contributes to mycobacterial virulence and, possibly, new vulnerabilities for chemotherapeutic and immunological interventions.

## Materials and Methods

### Bacterial strains and media

*M*.*tb* CDC1551 WT and Δesx1 mutant strains were cultured in 7H9 broth supplemented with 0.5% glycerol, 10% OADC, and 0.05% Tween 80 (henceforth ‘complete 7H9’). The Δesx5 mutant strain was cultured in a 1:1 mixture of complete 7H9 and a previously described nutrient-replete rich medium termed ‘YM medium’ (61). For any comparison between the Δesx5 mutant and other strains, all strains were maintained in identical media for a minimum of 48 hr. Liquid cultures were grown at 37°C with shaking at 180– 200 rpm. Bacterial growth plates were composed of 7H11 agar supplemented with 0.5% glycerol and 10% OADC. Induction medium was prepared by supplementing 7H9 broth with 0.1% glycerol, 1 mM sodium propionate, and 100 mM 2-(4-morpholino)-ethane sulfonic acid (MES) and adjusting to pH 6.5, adapted from (40). For lipid extraction, a modified 7H9 broth containing 0.5% glycerol, 10% OADC, 0.1 mM sodium propionate, and 0.05% Tyloxapol was used to facilitate virulence lipid formation, as demonstrated by (42). Selection of mutant strains was maintained using 50 ug/mL hygromycin.

### *M*.*tb* deletion strain construction

Specialized transduction phages carrying barcode sequences for deleting the ESX-5 and ESX-1 operons of *M*.*tb* were constructed as described previously (37). The details of primers used for amplifying the flanking regions and confirming the deletion mutants are included in **Table S1**. The primer combinations for three primer PCR verification of mutants are ESX-1_PCR_1, ESX-1_PCR_2, and Uptag for the Δesx1 strain and ESX-5_PCR_1, ESX-5_PCR_2, and Uptag for the Δesx5 strain. Both Δesx1 and Δesx5 strains were further confirmed by whole genome sequencing (**Fig. S1**). Genomic DNA isolated from these mutants were sequenced using NextseqXplus (2×151bp PE) platform (Illumina, USA), and the trimmed reads were mapped to the *M*.*tb* CDC1551 genome (NC_002755.2) using Bowtie2 (v 7.2.2), and the deletions were verified.

### *M*.*tb* protein isolation

To extract culture filtrate (CF) and outer mycomembrane (OMM) fractions from *M*.*tb* cultures, 50 mL bacterial cultures were grown to the mid-log phase (defined by OD_600_ 0.6). Bacteria were washed twice with tris-buffered saline (TBS), transferred into 50 mL induction medium, and incubated for 3 days at 37°C with shaking. Following induction, bacteria were pelleted, and the CF was transferred to a clean tube. The bacteria pellet was washed twice with TBS and resuspended in 500 μL of 1% OBG (Sigma) in TBS. Protease Inhibitor Cocktail for bacterial cell extracts (Sigma) at 1:100 dilution and 1 mM phenylmethylsulfonyl fluoride (PMSF) (Sigma) were added to the CF and the OBG solution. CF was concentrated by centrifugation through 20 mL 3 kDa molecular weight cuff-off spin columns (Cytiva) at 4°C to a final volume of ≤1 mL. Bacteria were incubated at 37°C for 30 min in the OBG solution, then pelleted by centrifugation, and the soluble fraction was harvested.

### Detergent susceptibility assay

To assess whether treatment with OBG caused lysis of *M*.*tb*, four independent WT and Δesx5 mutant cultures were incubated in induction medium as described above. 1 mL of each induced culture was pelleted and resuspended in 200 uL of 1% OBG in TBS for 30 min at 37°C, while another 1 mL was incubated in 200 uL pure TBS under the same conditions as a control. Following incubation, cultures were diluted 1:5 in PBS, then plated onto 7H11 agar as a series of 1:10 dilutions. WT *M*.*tb* CFUs were counted at Week 3 to compare bacterial viability with and without OBG treatment, while Δesx5 CFUs were counted at Week 6.

### Western blotting

CF and OMM fractions were mixed with 4X Laemmli sample buffer (Bio-Rad) and boiled for 10 min at 95°C. Protein samples were loaded alongside PageRuler Plus Prestained Protein Ladder (Thermo) onto 4-15% Mini-PROTEAN TGX gels (Bio-Rad) for electrophoresis at 100 V for 1 hr. Transfer to a 0.22 um PVDF membrane (Bio-Rad) was achieved using a wet transfer protocol on ice at 80 V for 45 min. Membranes were blocked for 1 hr at room temperature in 5% milk blocking buffer in TBS-T (TBS + 0.1% Tween 20). Primary antibodies were incubated overnight at 4°C in 5% milk, followed by washing with TBS-T. Secondary antibodies were incubated for 1 hr at room temperature in TBS-T, followed by washing in TBS-T. Membranes were incubated with SuperSignal West Pico PLUS substrate (Thermo) and imaged with a Kwik Quant imager (Kindle Biosciences).

Anti-PE_PGRS33 (7C4.1F7) primary was obtained from the International AIDS Vaccine Initiative and was originally developed in (19). Note that this anti-PE_PGRS33 antibody has been previously shown to react with multiple PE_PGRS family proteins, presumably due to high similarity (26, 35). Anti-Ag85 primary was obtained through BEI Resources, NIAID, NIH as follows: Monoclonal Anti-Mycobacterium tuberculosis Ag85 Complex (FbpA/FbpB/FbpC; Genes Rv3804c, Rv1886c, Rv0129c), Clone CS-90 (produced *in vitro*), NR-13816. Anti-EsxB primary was obtained through BEI Resources, NIAID, NIH as follows: Polyclonal Anti-Mycobacterium tuberculosis CFP10 (Gene Rv3874) (antiserum, Rabbit), NR-13801. Anti-rabbit and anti-mouse secondary antibodies were purchased from Cell Signaling Technologies (#7074 and #7076, respectively).

### Mass spectrometry

CF and OMM fractions from *M*.*tb* were obtained as above. Subsequently, samples were reduced with tris(2-carboxyethyl)phosphine (TCEP) (Thermo), alkylated with iodoacetamide (MP Biochemicals), and further reduced with dithiothreitol (DTT) (Alfa Aesar). Samples were digested with Lys-C (Wako) overnight at room temperature and trypsin (Thermo) for 6 hr at 37°C. Samples were labeled with TMT 6-plex reagents (Thermo) and pooled by normalizing to the amount of labeled sample. Peptides were desalted by STAGE tips and analyzed on an Orbitrap Eclipse mass spectrometer (Thermo) with a FAIMS device enabled. MS2 spectra were searched against an *M*.*tb* Uniprot composite database using the COMET tool, and peptide-spectrum matches were filtered to a false-discovery rate of 1% (62). Peptide abundances were normalized to the total peptide abundance for each sample. Functional annotation analysis was conducted by inputting lists of differentially expressed genes (≥2-fold differential expression, p < 0.05) into the DAVID functional annotation tool (63, 64).

### *M*.*tb* RNA isolation

Total RNA from WT and Δesx5 cultures were isolated using a Direct-zol RNA Miniprep plus kit (Zymo Research, USA). Briefly, 10 ml cultures in duplicate were grown to mid-log phase in a 1:1 mixture of complete 7H9 and YM medium. The cells were washed twice using phosphate-buffered saline (PBS) and incubated in induction medium for 3 days at 37°C, with shaking. The cell pellets were resuspended in 1 ml of RNAlater (Qiagen, USA) and stored overnight at 4°C. RNA isolation was performed following the manufacturer’s instructions, and the total RNA was quantified using Qubit RNA BR Assay Kits (ThermoFischer, USA).

### RNA-seq

Following total RNA extraction, the genomic libraries were prepared using Zymo-Seq Ribofree Total RNA Library kit (Zymo Research, USA) following the manufacturer’s instructions. This kit included cDNA synthesis, ribosomal RNA depletion, and library preparation reagents. The genomic libraries were sequenced using Illumina NextSeq 2000 platform using a P1 reagents kit (150 PE). Sequencing was performed in duplicate for both WT and Δesx5. RNA-seq reads were analyzed using Geneious Prime (v 2024.0.2). BBDuk trimmer (Biomatters ltd.), and FastQC (v0.11.9) were used to trim and perform quality control on the reads. Processed reads were mapped to the *M*.*tb* CDC1551 genome (NC_002755.2) using Bowtie2 (v 7.2.2). The RPKM (Reads per kilobase per million), TPM (Transcripts per million) and FPKM (Fragments Per Kilobase per Million mapped reads) were counted. The differential expression analysis was performed using DeSeq2, and expression clustering was calculated using log2FoldChange. Raw RNA-seq reads are available at the Sequence Read Archive (SRA) database at accession number PRJNA1246567. Individual samples can be accessed as follows: SAMN47784052 (WT_1), SAMN47784053 (WT_2), SAMN47784054 (Δesx5_1), SAMN47784055 (Δesx5_2).

### *M*.*tb* lipid extraction

For lipid extraction, 20 mL *M*.*tb* cultures were grown to OD_600_ 1.5. To label methyl-branched lipids, 1 μCi/mL ^14^C-propionate was added to each culture, and samples were incubated for 24 hr at 37°C with shaking. Extraction of cell surface lipids was performed as previously described (65, 66). Briefly, ^14^C-propionate labeled bacteria were incubated with 1:2 chloroform:methanol for 24 hr followed by 2:1 chloroform:methanol for 48 hr to extract cell surface lipids. Organic fractions were washed twice with distilled water and dried under airflow. Samples were resuspended in a volume of dichloromethane proportionate to sample mass, and radiolabel uptake was measured using a Beckman LS 6000SE liquid scintillation counter.

### Thin-layer chromatography

Running solvents for TLC were adapted from (65). Briefly, 100,000 cpm of each sample was loaded onto aluminum-backed TLC plates and allowed to run in one direction. Cell surface lipid fractions were run using 98:2 petroleum ether:acetone solvent. TLC plates were exposed to a phosphorimaging cassette for 24 hours and visualized on an Amersham Typhoon RGB phosphorimager (Cytiva). Background correction and semiquantitative band detection was conducted using ImageQuant TL software. To measure pixel intensities across each lane of the TLC plate, the plot profile function in ImageJ was used. The mean background pixel intensity was determined from the portion of the image surrounding the lanes and was subtracted from the intensity values.

### Scanning electron microscopy

To image the bacterial surface, *M*.*tb* strains were grown to OD_600_ 1.5. Cultures were washed into an equal volume of 50:50 complete 7H9:YM medium or induction medium, then 1 mL aliquots were pipetted atop poly-L-lysine treated glass coverslips in a 24-well plate. 6 mM DTT (Sigma) was added to each well to promote *M*.*tb* adhesion to the coverslip, adapted from (67). Bacteria were incubated for 48 hr at 37°C without shaking then washed once with PBS. Coverslips were fixed in 4% paraformaldehyde (Electron Microscopy Sciences) plus 2% glutaraldehyde (Electron Microscopy Sciences) in PBS overnight at room temperature. Samples were post-fixed in 1% osmium tetroxide in 0.1 M sodium cacodylate for 1 hr on ice in the dark. Samples were rinsed with distilled water then dehydrated in a graded ethanol series and left to dry overnight in a desiccator with hexamethyldisilazane (HMDS). Samples were mounted on carbon coated stubs, coated with 5 nm gold-palladium (Denton DeskV), and imaged on a JEOL SEM (JSM-IT700HR) at 3 kV.

### Antibiotic susceptibility testing

To assess the susceptibility of *M*.*tb* WT and Δesx5 to antibiotic compounds, a microplate-based alamarBlue assay was used as described previously (47, 48). Briefly, a series of 2-fold dilutions of stock solutions of each compound in complete 7H9 were prepared. Bacteria were grown to mid-log OD and diluted in complete 7H9 to a concentration of 10^5^ CFU per 100 uL in each well. To each strain, 100 uL of each antibiotic dilution was added. Plates were incubated at 37°C for 7 days for WT *M*.*tb* or 10 days for Δesx5; then fluorescence intensity was measured on a microplate reader using an excitation of 544 nm and an emission of 590 nm. Percent inhibition was computed by comparing relative fluorescence to the untreated control for each respective strain for each plate. MIC90 (the minimum concentration required to achieve ≥90% inhibition) was used to define MIC.

## Data availability

Raw proteomic data for CF and OMM fractions have been provided as Microsoft Excel files, **Dataset S1** and **Dataset S2**, respectively. Raw transcriptomic data have been provided as a Microsoft Excel file, **Dataset S3**, and raw data is provided at the accension numbers provided in Methods. All other data generated as part of this study are available upon request to the corresponding author.

## Acknowledgments

We thank the members of the Thermo Fisher Scientific Center for Multiplexed Proteomics at Harvard Medical School for conducting the mass spectrometry quantification. We also thank Barbara Smith of the Johns Hopkins University School of Medicine’s Microscope Facility for performing the scanning electron microscopy imaging.

The authors gratefully acknowledge the support of NIH grants R01AI152688 (WRB), R01AI155602 (WRB), R37AI167750 (WRB), R01AI026170 (WRJ), and R24AI134650 (WRJ).

## References

1. S. M. Gygli, S. Borrell, A. Trauner, S. Gagneux, Antimicrobial resistance in Mycobacterium tuberculosis: mechanistic and evolutionary perspectives. FEMS Microbiol Rev 41, 354–373 (2017).

2. M. F. Goldberg, N. K. Saini, S. A. Porcelli, Evasion of Innate and Adaptive Immunity by Mycobacterium tuberculosis. Microbiol Spectr 2 (2014).

3. M. B. Goren, P. J. Brennan, “Tuberculosis” in Mycobacterial Lipids: Chemistry and Biologic Activities, G. P. Youmans, Ed. (W.B. Saunders Company, Philadelphia, PA, 1979), pp. 63–193.

4. M. Daffé, P. Draper, The envelope layers of mycobacteria with reference to their pathogenicity. Adv Microb Physiol 39, 131–203 (1998).

5. T. A. Sysoeva, M. A. Zepeda-Rivera, L. A. Huppert, B. M. Burton, Dimer recognition and secretion by the ESX secretion system in Bacillus subtilis. Proc Natl Acad Sci U S A 111, 7653–7658 (2014).

6. C. Poulsen, S. Panjikar, S. J. Holton, M. Wilmanns, Y. H. Song, WXG100 protein superfamily consists of three subfamilies and exhibits an α-helical C-terminal conserved residue pattern. PLoS One 9, e89313 (2014).

7. A. S. Pym, P. Brodin, R. Brosch, M. Huerre, S. T. Cole, Loss of RD1 contributed to the attenuation of the live tuberculosis vaccines Mycobacterium bovis BCG and Mycobacterium microti. Mol Microbiol 46, 709–717 (2002).

8. T. Hsu et al., The primary mechanism of attenuation of bacillus Calmette-Guerin is a loss of secreted lytic function required for invasion of lung interstitial tissue. Proc Natl Acad Sci U S A 100, 12420–12425 (2003).

9. M. S. Siegrist et al., Mycobacterial Esx-3 is required for mycobactin-mediated iron acquisition. Proc Natl Acad Sci U S A 106, 18792–18797 (2009).

10. A. Serafini, D. Pisu, G. Palù, G. M. Rodriguez, R. Manganelli, The ESX-3 secretion system is necessary for iron and zinc homeostasis in Mycobacterium tuberculosis. PLoS One 8, e78351 (2013).

11. V. Boradia, A. Frando, C. Grundner, The Mycobacterium tuberculosis PE15/PPE20 complex transports calcium across the outer membrane. PLoS Biol 20, e3001906 (2022).

12. D. Pajuelo et al., Toxin secretion and trafficking by Mycobacterium tuberculosis. Nat Commun 12, 6592 (2021).

13. N. Sankey et al., Role of the Mycobacterium tuberculosis ESX-4 Secretion System in Heme Iron Utilization and Pore Formation by PPE Proteins. mSphere 8, e0057322 (2023).

14. R. R. Nair, V. Meikle, S. Dubey, M. Pavlenok, M. Niederweis, Master control of protein secretion by Mycobacterium tuberculosis. bioRxiv (2025).

15. N. C. Gey van Pittius et al., Evolution and expansion of the Mycobacterium tuberculosis PE and PPE multigene families and their association with the duplication of the ESAT-6 (esx) gene cluster regions. BMC Evol Biol 6, 95 (2006).

16. S. T. Cole et al., Deciphering the biology of Mycobacterium tuberculosis from the complete genome sequence. Nature 393, 537–544 (1998).

17. S. B. Snapper et al., Lysogeny and transformation in mycobacteria: stable expression of foreign genes. Proc Natl Acad Sci U S A 85, 6987–6991 (1988).

18. S. B. Snapper, R. E. Melton, S. Mustafa, T. Kieser, W. R. Jacobs, Isolation and characterization of efficient plasmid transformation mutants of Mycobacterium smegmatis. Mol Microbiol 4, 1911–1919 (1990).

19. A. M. Abdallah et al., PPE and PE_PGRS proteins of Mycobacterium marinum are transported via the type VII secretion system ESX-5. Mol Microbiol 73, 329–340 (2009).

20. D. Bottai et al., Disruption of the ESX-5 system of Mycobacterium tuberculosis causes loss of PPE protein secretion, reduction of cell wall integrity and strong attenuation. Mol Microbiol 83, 1195–1209 (2012).

21. A. Palande et al., Proteomic Analysis of the Mycobacterium tuberculosis Outer Membrane for Potential Implications in Uptake of Small Molecules. ACS Infect Dis 10, 890–906 (2024).

22. C. M. Sassetti, D. H. Boyd, E. J. Rubin, Genes required for mycobacterial growth defined by high density mutagenesis. Mol Microbiol 48, 77–84 (2003).

23. M. A. DeJesus et al., Comprehensive Essentiality Analysis of the Mycobacterium tuberculosis Genome via Saturating Transposon Mutagenesis. mBio 8 (2017).

24. N. A. Kruh, J. Troudt, A. Izzo, J. Prenni, K. M. Dobos, Portrait of a pathogen: the Mycobacterium tuberculosis proteome in vivo. PLoS One 5, e13938 (2010).

25. J. Qian, R. Chen, H. Wang, X. Zhang, Role of the PE/PPE Family in Host-Pathogen Interactions and Prospects for Anti-Tuberculosis Vaccine and Diagnostic Tool Design. Front Cell Infect Microbiol 10, 594288 (2020).

26. L. S. Ates et al., RD5-mediated lack of PE_PGRS and PPE-MPTR export in BCG vaccine strains results in strong reduction of antigenic repertoire but little impact on protection. PLoS Pathog 14, e1007139 (2018).

27. T. P. Stinear et al., Insights from the complete genome sequence of Mycobacterium marinum on the evolution of Mycobacterium tuberculosis. Genome Res 18, 729–741 (2008).

28. F. Guo et al., Immunological effects of the PE/PPE family proteins of Mycobacterium tuberculosis and related vaccines. Front Immunol 14, 1255920 (2023).

29. Q. Wang et al., PE/PPE proteins mediate nutrient transport across the outer membrane of Mycobacterium tuberculosis. Science 367, 1147–1151 (2020).

30. M. Di Luca et al., The ESX-5 associated eccB-EccC locus is essential for Mycobacterium tuberculosis viability. PLoS One 7, e52059 (2012).

31. L. S. Ates et al., Essential Role of the ESX-5 Secretion System in Outer Membrane Permeability of Pathogenic Mycobacteria. PLoS Genet 11, e1005190 (2015).

32. E. M. Weerdenburg et al., ESX-5-deficient Mycobacterium marinum is hypervirulent in adult zebrafish. Cell Microbiol 14, 728–739 (2012).

33. D. W. White, S. R. Elliott, E. Odean, L. T. Bemis, A. D. Tischler, Pst/SenX3-RegX3 Regulates Membrane Vesicle Production Independently of ESX-5 Activity. mBio 9 (2018).

34. S. Tiwari et al., BCG-Prime and boost with Esx-5 secretion system deletion mutant leads to better protection against clinical strains of Mycobacterium tuberculosis. Vaccine 38, 7156–7165 (2020).

35. L. S. Ates et al., Mutations in ppe38 block PE_PGRS secretion and increase virulence of Mycobacterium tuberculosis. Nat Microbiol 3, 181–188 (2018).

36. J. Gallant et al., PPE38-Secretion-Dependent Proteins of M. tuberculosis Alter NF-kB Signalling and Inflammatory Responses in Macrophages. Front Immunol 12, 702359 (2021).

37. P. Jain et al., Specialized transduction designed for precise high-throughput unmarked deletions in Mycobacterium tuberculosis. mBio 5, e01245–01214 (2014).

38. C. Hoffmann, A. Leis, M. Niederweis, J. M. Plitzko, H. Engelhardt, Disclosure of the mycobacterial outer membrane: cryo-electron tomography and vitreous sections reveal the lipid bilayer structure. Proc Natl Acad Sci U S A 105, 3963–3967 (2008).

39. A. D. van der Woude et al., Differential detergent extraction of mycobacterium marinum cell envelope proteins identifies an extensively modified threonine-rich outer membrane protein with channel activity. J Bacteriol 195, 2050–2059 (2013).

40. M. E. Feltcher et al., Label-free Quantitative Proteomics Reveals a Role for the Mycobacterium tuberculosis SecA2 Pathway in Exporting Solute Binding Proteins and Mce Transporters to the Cell Wall. Mol Cell Proteomics 14, 1501–1516 (2015).

41. G. Leisching, R. D. Pietersen, I. Wiid, B. Baker, Virulence, biochemistry, morphology and host-interacting properties of detergent-free cultured mycobacteria: An update. Tuberculosis (Edinb) 100, 53–60 (2016).

42. C. V. Mulholland et al., Propionate prevents loss of the PDIM virulence lipid in Mycobacterium tuberculosis. Nat Microbiol 9, 1607–1618 (2024).

43. J. M. Tufariello et al., Separable roles for Mycobacterium tuberculosis ESX-3 effectors in iron acquisition and virulence. Proc Natl Acad Sci U S A 113, E348–357 (2016).

44. T. H. Phan, R. Ummels, W. Bitter, E. N. Houben, Identification of a substrate domain that determines system specificity in mycobacterial type VII secretion systems. Sci Rep 7, 42704 (2017).

45. T. Mori et al., Specific detection of tuberculosis infection: an interferon-gamma-based assay using new antigens. Am J Respir Crit Care Med 170, 59–64 (2004).

46. M. Marrichi, L. Camacho, D. G. Russell, M. P. DeLisa, Genetic toggling of alkaline phosphatase folding reveals signal peptides for all major modes of transport across the inner membrane of bacteria. J Biol Chem 283, 35223–35235 (2008).

47. L. Collins, S. G. Franzblau, Microplate alamar blue assay versus BACTEC 460 system for high-throughput screening of compounds against Mycobacterium tuberculosis and Mycobacterium avium. Antimicrob Agents Chemother 41, 1004–1009 (1997).

48. S. Lun et al., Indoleamides are active against drug-resistant Mycobacterium tuberculosis. Nat Commun 4, 2907 (2013).

49. O. T. Schubert et al., The Mtb proteome library: a resource of assays to quantify the complete proteome of Mycobacterium tuberculosis. Cell Host Microbe 13, 602–612 (2013).

50. L. S. Ates, New insights into the mycobacterial PE and PPE proteins provide a framework for future research. Mol Microbiol 113, 4–21 (2020).

51. S. Fishbein, N. van Wyk, R. M. Warren, S. L. Sampson, Phylogeny to function: PE/PPE protein evolution and impact on Mycobacterium tuberculosis pathogenicity. Mol Microbiol 96, 901–916 (2015).

52. S. R. Elliott, A. D. Tischler, Phosphate starvation: a novel signal that triggers ESX-5 secretion in Mycobacterium tuberculosis. Mol Microbiol 100, 510–526 (2016).

53. R. Copin et al., Sequence Diversity in the pe_pgrs Genes of Mycobacterium tuberculosis Is Independent of Human T Cell Recognition. mBio 5, e00960–e00913 (2014).

54. T. Sharma et al., The Mycobacterium tuberculosis PE_PGRS Protein Family Acts as an Immunological Decoy to Subvert Host Immune Response. Int J Mol Sci 23 (2022).

55. B. R. Gordon et al., Lsr2 is a nucleoid-associated protein that targets AT-rich sequences and virulence genes in Mycobacterium tuberculosis. Proc Natl Acad Sci U S A 107, 5154–5159 (2010).

56. O. S. Rosenberg et al., EspR, a key regulator of Mycobacterium tuberculosis virulence, adopts a unique dimeric structure among helix-turn-helix proteins. Proc Natl Acad Sci U S A 108, 13450–13455 (2011).

57. D. M. Hunt et al., Long-range transcriptional control of an operon necessary for virulence-critical ESX-1 secretion in Mycobacterium tuberculosis. J Bacteriol 194, 2307–2320 (2012).

58. A. Garces et al., EspA acts as a critical mediator of ESX1-dependent virulence in Mycobacterium tuberculosis by affecting bacterial cell wall integrity. PLoS Pathog 6, e1000957 (2010).

59. L. S. Ates et al., The ESX-5 System of Pathogenic Mycobacteria Is Involved In Capsule Integrity and Virulence through Its Substrate PPE10. PLoS Pathog 12, e1005696 (2016).

60. C. Mailaender et al., The MspA porin promotes growth and increases antibiotic susceptibility of both Mycobacterium bovis BCG and Mycobacterium tuberculosis. Microbiology (Reading) 150, 853–864 (2004).

61. Y. Minato et al., Genomewide Assessment of Mycobacterium tuberculosis Conditionally Essential Metabolic Pathways. mSystems 4 (2019).

62. J. K. Eng, T. A. Jahan, M. R. Hoopmann, Comet: an open-source MS/MS sequence database search tool. Proteomics 13, 22–24 (2013).

63. d. W. Huang, B. T. Sherman, R. A. Lempicki, Systematic and integrative analysis of large gene lists using DAVID bioinformatics resources. Nat Protoc 4, 44–57 (2009).

64. B. T. Sherman et al., DAVID: a web server for functional enrichment analysis and functional annotation of gene lists (2021 update). Nucleic Acids Res 50, W216–W221 (2022).

65. A. Singh et al., Mycobacterium tuberculosis WhiB3 maintains redox homeostasis by regulating virulence lipid anabolism to modulate macrophage response. PLoS Pathog 5, e1000545 (2009).

66. S. Shee et al., Biosensor-integrated transposon mutagenesis reveals rv0158 as a coordinator of redox homeostasis in Mycobacterium tuberculosis. Elife 12 (2023).

67. A. Trivedi, P. S. Mavi, D. Bhatt, A. Kumar, Thiol reductive stress induces cellulose-anchored biofilm formation in Mycobacterium tuberculosis. Nat Commun 7, 11392 (2016).

