## Supplementary Figures for "Loss of the ESX-5 secretion locus in *Mycobacterium tuberculosis* reshapes the mycomembrane and enhances ESX-1 substrate secretion"

**This PDF file includes:**

Figures S1 to S4

**Other supporting materials for this manuscript include the following:**

Table S1
Datasets S1 to S3

### Supplementary Figures

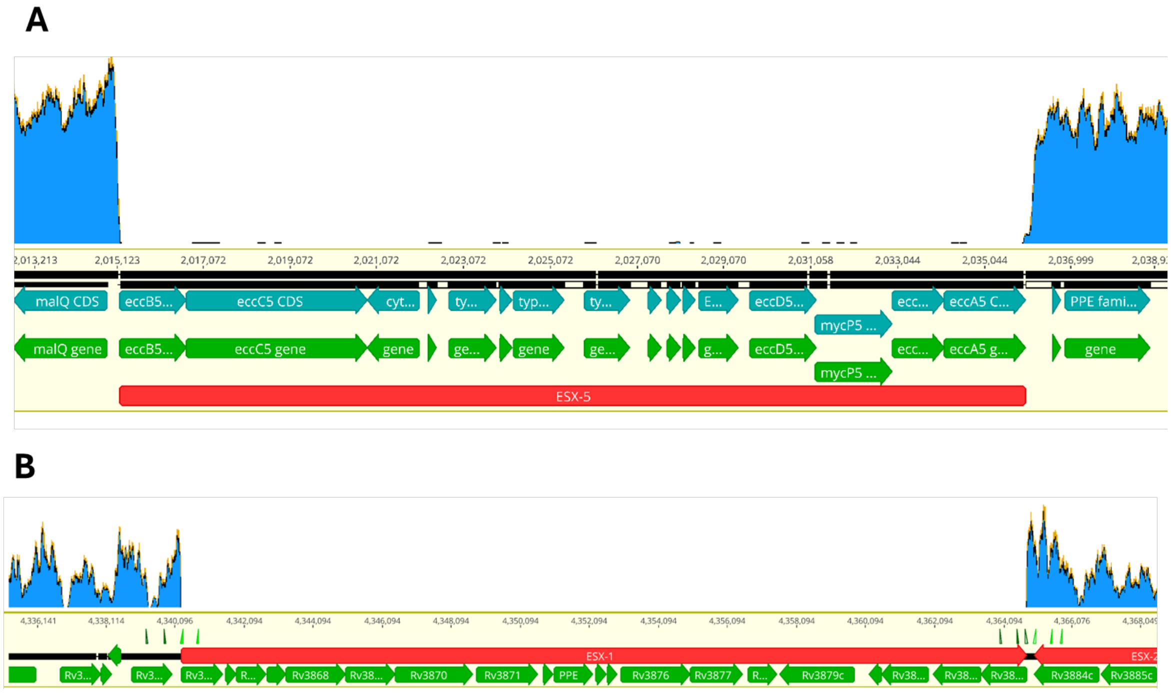


**Figure S1: Sequence confirmation of *M.tb* ∆esx5 and ∆esx1 mutant strains.** (A–B) Barcoded deletion mutant strains ∆esx5 and ∆esx1 were generated in an *M.tb* CDC1551 background by specialized transduction. Deletions comprising the 17 genes in the ESX-5 operon (A) and the 20 genes in the ESX-1 operon (B) were confirmed by Illumina short-read (150 bp paired-end) whole genome sequencing, which revealed a complete absence of reads mapping to the deleted regions.

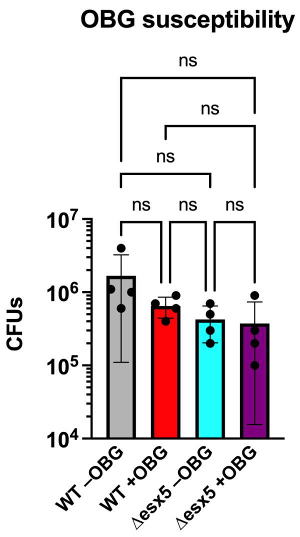


**Figure S2: *M.tb* treatment with OBG detergent is non-lytic.**
*M.tb* CDC1551 WT and Δesx5 mutant bacteria were incubated in tris-buffered saline containing either no detergent or 1% OBG for 30 min at 37°C. The resulting cells were plated onto 7H11, and viability was assessed by CFU enumeration at either Week 3 (for WT) or Week 6 (for Δesx5). No significant difference between any strain or treatment condition was noted by one-way ANOVA. Error bars reflect mean ± standard deviation (n=4 each, ns: non-significant).

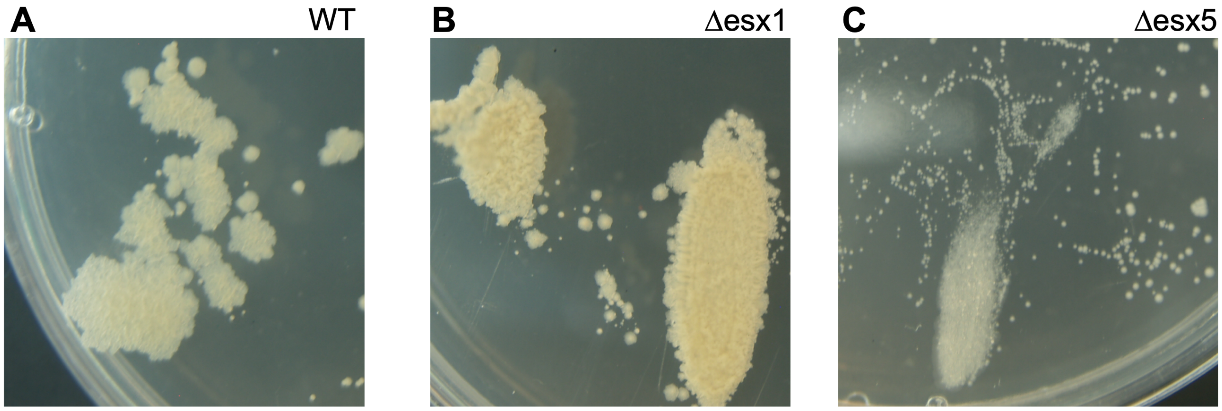


**Figure S3: The *M.tb* ∆esx5 mutant shows altered colony morphology.** (A–C) Images of WT (A), ∆esx1 (B), and ∆esx5 (C) *M.tb* strains grown on 7H11 agar. Mid-log phase cultures were swabbed onto agar using 10 uL inoculation loops and incubated at 37°C for 6 weeks. Compared to WT and the ∆esx1 mutant, the ∆esx5 mutant yielded smaller colonies with a smoother surface appearance.

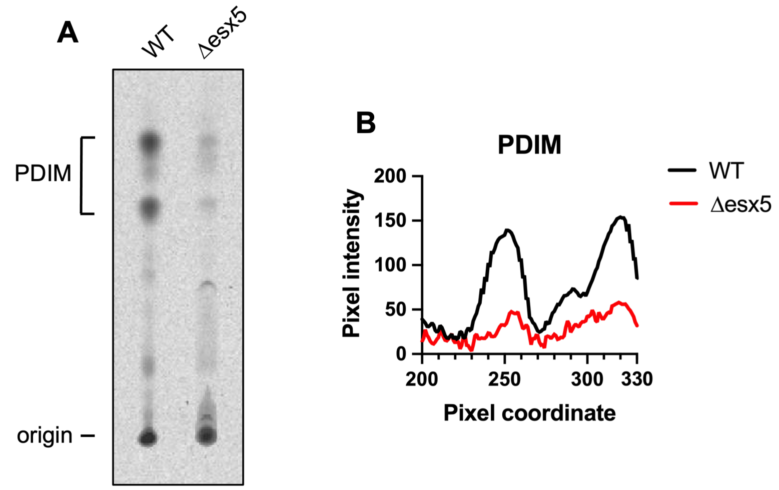


**Figure S4: The *M.tb* Δesx5 mutant produces less PDIM virulence lipids.**
(A) Total lipids from *M.tb* WT and Δesx5 were labeled by addition of $^{14}C$-propionate. Polar lipids were extracted, and 100,000 cpm of each sample were loaded onto a TLC plate. Plates were developed in 98:2 petroleum ether:acetone and visualized by phosphorimaging. Locations of phthiocerol dimycocerosate (PDIM) lipids and the origin are indicated. TLC plate reflects one replicate of three independent experiments performed from different bacteria cultures.
(B) Pixel intensities (from 0–255) were measured along the lanes of the TLC plate, beginning at the origin. WT and the Δesx5 mutant feature similarly shaped peaks in the region corresponding to PDIM lipids, although the intensity is lower for the Δesx5 mutant.

### Supplementary Tables

**Table S1: Primers used in this study.**
*(Please see attached Excel file.)*

### Supplementary Datasets

**Dataset S1: Mass spectrometry outputs for the culture filtrate fraction of *M.tb* WT and ∆esx5 strains.**
Raw peptides and quantified proteins, normalized to total peptides per sample by summed signal/noise (S/N) ratio, are provided. $Log_2$ fold-changes and p-values were computed for each comparison.

**Dataset S2: Mass spectrometry outputs for the outer mycomembrane fraction of *M.tb* WT and ∆esx5 strains.**
Raw peptides and quantified proteins, normalized to total peptides per sample by summed signal/noise (S/N) ratio, are provided. $Log_2$ fold-changes and p-values were computed for each comparison.

**Dataset S3: RNAseq output for transcripts from *M.tb* WT and ∆esx5 strains.**
Differential expression values for the ∆esx5 / WT comparison, normalized to read depth, are provided. $Log_2$ fold-changes and p-values were computed for each comparison.
